# AI-driven analysis for real-time detection of unstained microscopic cell culture images

**DOI:** 10.1101/2025.07.27.667077

**Authors:** Kathrin Hildebrand, Tatiana Mögele, Dennis Raith, Maria Kling, Anna Rubeck, Stefan Schiele, Eelco Meerdink, Avani Sapre, Jonas Bermeitinger, Martin Trepel, Rainer Claus

**Author notes:** Corresponding author: Rainer Claus, Adress: Stenglinstr. 2, 86156 Augsburg, Germany, Fon. both first authors contributed equally (Kathrin Hildebrand, Tatiana Mögele).

## Abstract

AI-based image recognition has significantly advanced the analysis of tissues and individual cells both in the context of translational studies and diagnostics. To date, recognition is primarily based on the identification of certain cell characteristics (e.g. by staining). The morphological assessment of unstained cells holds additional potential, as it allows for virtually real-time assessment without the need to manipulate the cells. This facilitates longitudinal observations, as required for drug testing, and forms a basis for autonomous experimental execution.

A semi-automated cell culture system (AICE3, LabMaite) was used to culture myeloid leukemic cell lines (K562, HL-60, Kasumi-1). K562 cells were treated with hemin and PMA to induce erythroid and megakaryocytic differentiation, respectively. Cell images were acquired using automated bright field microscopy. Images were used to train an AI model using an NVIDIA DGX A100 GPU with Ultralytics YOLOv8. Morphologic features were extracted using RedTell.

The model reliably distinguished K562 cells from HL-60 and Kasumi-1 using >400 images per class (average >15 cells/image). Bounding boxes were generated correctly (mAP@.5 >98%); precision and sensitivity exceeded 97%. Validation on an external K562 dataset confirmed these results. Classification of all three cell lines achieved >97% sensitivity/specificity and 94.6% precision.

To test drug response, we used YOLOv8-s to distinguish untreated K562 cells from those undergoing erythroid or megakaryocytic differentiation (n >3,000 annotations). Precision, sensitivity, and specificity were >95%. RedTell identified 3 of 74 morphological traits contributing significantly to class separation.

We demonstrate accurate, near real-time detection of unstained cells, enabling future AI-based drug testing.

## Introduction

In the search for new cancer therapeutics, in vitro drug testing and drug screening in cell lines and on primary samples and their derivatives have proven to be indispensable tools that accelerate the drug discovery process. This approach allows researchers to evaluate the efficacy and safety of potential drug candidates before advancing to more complex and ethically challenging *in vivo* studies (1–4). One of the primary advantages of *in vitro* drug testing is the ability to conduct experiments in a highly controlled environment. Cell lines provide a simplified yet representative model of cellular behavior. This enables precise investigation of the impact of drug candidates on phenotypic changes and specific molecular pathways. Furthermore, it facilitates high-throughput screening, allowing to assess a large number of compounds efficiently and accelerating the drug discovery timeline by quickly identifying promising candidates for further investigation. As such, *in vitro* drug testing may have utmost relevance in personalized medicine allowing researchers to tailor drug-screening assays to individual genetic profiles, predicting how a patient might respond to a specific treatment and optimizing therapeutic strategies (5). Automated and advanced imaging technologies further enhance the speed and accuracy of compound screening, enabling researchers to sift through vast libraries of potential drugs and combinations.(6)

However, the ultimate utility of cellular models for drug testing depends on accurate real-time longitudinal monitoring and recognizing relevant molecular and/or phenotypic changes such as differentiation, induction of cell death and others (7). Molecular analyses at different omics levels (e.g. genetic profiling) and functional readouts are the basis for characterizing the cellular response to different stimuli. Staining, e. g. by immunofluorescence, can detect and characterize cellular responses but relies on the expression of specific cell surface antigens and prevents “untouched” longitudinal observation. Also, these target markers may not be available or specific enough, samples may be too heterogeneous and the specific labeling may affect the cell, e.g. by activation or loss of viability (8). Thus, assessment of cell morphology from unstained cell culture images might overcome the limitations of staining techniques with exogenous dyes, enabling the observation of cells in their native state and offering insights into cellular dynamics and behavior in real-time.

Computer-aided approaches based on image processing enable fast, high-throughput and reliable assessment of cell morphology while minimizing deviations from human observation. The development of automatic microscopes has enabled the creation of large image datasets, which have been extensively studied using computational tools. Deep learning (DL) techniques have been successfully used to automatically classify and segment images and various cell types, including motor neurons, stem cells, and others, have been accurately classified using deep convolutional neural networks (CNN) trained with labeled images (9–11). Previous machine learning methods for image-based cell phenotyping have been usually based on the manual selection of individual features and image characteristics (such as size, shape, texturing of the cytoplasm and nucleus) (9, 12). However, the ability to identify specific phenotypes is limited by the diversity and complexity of the features, which in turn limits the ability to recognize and evaluate subtle changes and e.g. specific responses to certain substances. Therefore, previous studies have focused on large distinct cell groups such as lymphocytes, granulocytes and erythrocytes, whose morphology is easily recognizable to the human eye (12). Computer-assisted phenotyping of cells with more subtle charactersitics and morphological changes, as expected in drug testing, has not been established yet.

This study aims to demonstrate the promising potential of a real-time, image-based monitoring approach for detecting subtle morphological changes in unstained leukemia cells. Using DL and CNN, we utilize a semi-automated cell culture system with automated imaging to generate large datasets suitable for high-throughput, stain-free analysis. This approach allows continuous real-time evaluation of drug-induced morphological changes and presents a novel contribution towards “untouched” dynamic *in vitro* drug testing.

## Material and methods

### Cell culture and in vitro treatment

The myeloid cell lines K562, HL-60, and Kasumi-1 (German Collection of Microorganisms and Cell Cultures GmbH, DSMZ, Braunschweig, Germany) were cultured in RPMI1640 media (Gibco) containing 5% Penicillin/Streptomycin (Gibco) and 10% (K562) or 20% (HL-60, Kasumi-1) heat-inactivated fetal calf serum (Sigma-Aldrich). Cells were maintained at 0,2 x10^6^/cells per ml in T75 flasks and split every 3 to 4 days.

K562 cells were treated with 50 µM hemin (Sigma-Aldrich) for six days and with 5 nM phorbol myristate acetate (PMA, Sigma-Aldrich) (PMA) for 48 h for induction of erythroid differentiation and megakaryocytic differentiation, respectively. Decitabine (Sigma-Aldrich) treatment was conducted with three 20 or 100 nM pulses in 24 h intervals and cells were harvested after six days, as previously described by Stomper et al. (13). Cell numbers were counted by 0,4% trypan blue staining and viability was measured using the NucleoCounter system (Chemometec). The terms “Hemin”, “PMA” and “DAC” are used as abbreviations representing the cell population treated with the respective agent.

### Automated cell culture and image generation

For image generation cells were transferred to a 50 ml falcon (reactor) with a minimum concentration of 0.2 x10^6^ cells/ml in 20 ml media. All preparations of the reactor were carried out under a laminar flow. The reactor containing the cells was assembled for automated cell culture and image generation in an AICE3 device (LABMaiTE, Freiburg) inside of an incubator as described in Supplemental Figure 1. The device comprises a peristaltic pump system and a microscope with a 20x objective with an integrated camera and is kept in an incubator at 37°C and a 5% CO2 atmosphere to ensure the correct conditions for cell culture during imaging. To avoid pelleting in the reactor vial, the suspension was mixed every two minutes at a rate of 35 ml/min. At each step of image generation, five ml cell suspension were pumped from the reactor through a 0.2 µm Luer slide (ibidi GmbH, Gräfelfing, Germany) attached to the microscopic device. Cells were allowed to sediment on the slide for three minutes to ensure optimal focus before image generation. At each step, four different images were generated and cell counts were documented as the mean of the detected cells over all four images. With this setup, image generation was performed for 1.5 hours, resulting in 30 images per experimental run. Images without cells were used for background training. For longer experiments, half of the reactor was drained and refilled with fresh media about every 2,5 days and time between images was adapted to every 2 hours.

### Image processing and training

Roboflow (14) was used for annotation, splitting, and augmentation (flips, rotations). The annotation process was done manually for each cell type and segmentation masks were created using Roboflow’s smart polygon tool. The data was split into training, validation and test subsets (ratio 70:20:10) to facilitate effective model training, hyperparameter tuning, and performance evaluation and to ensure reliable and robust machine learning applications.

The models were trained on an NVIDIA DGX A100 using Ultralytics YOLOv8 environment (15). YOLOv8 is a single-shot state of the art object detection algorithm with a high accuracy rate on COCO (16) and Roboflow. Different sized YOLOv8-models (n, s, m, l) and Realtime Detection Transformers (RT-DETR) were taken into consideretation in training. The whole process from image generation to evaluation is depicted in Figure 1A. The code snippets used for the training, validation and prediction are given in Supplemental Figure 2, as well as the used confidence thresholds (objects with scores below this threshold were disregarded as false positives) and remaining hyperparameters (image size, training epochs, confidence threshold) (Supplemental Table 1).

**Figure 1:**
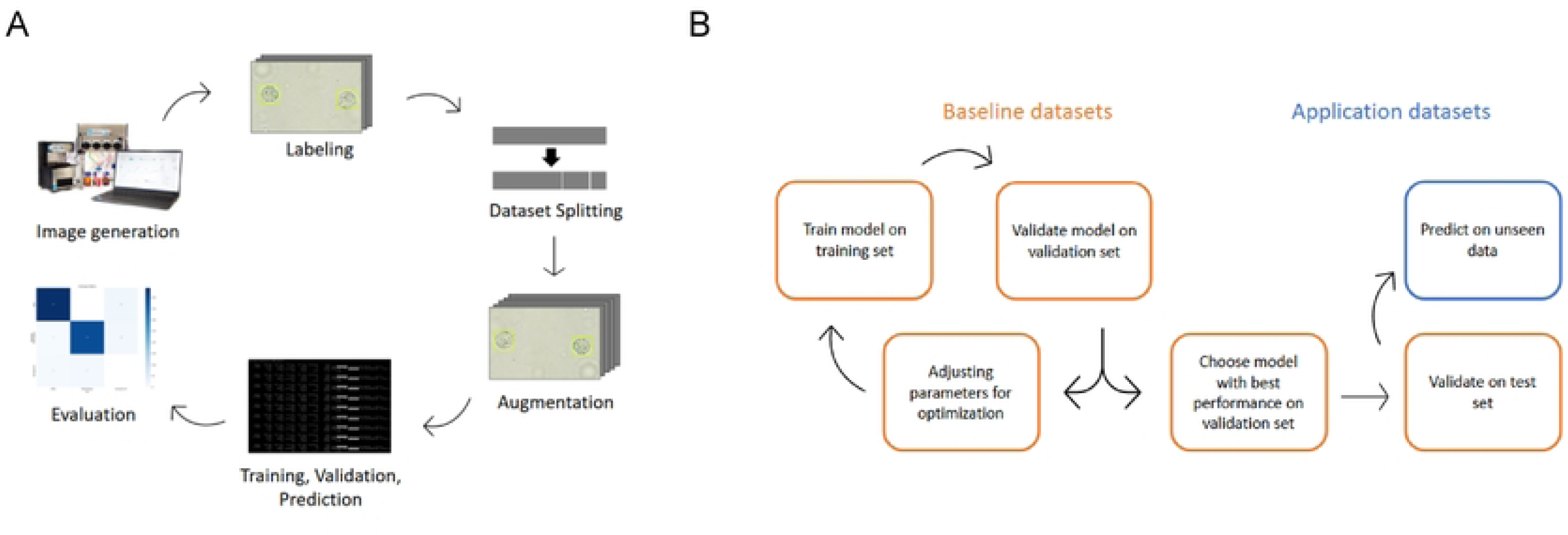
Overview of the deep learning workflow and model evaluation strategy. (A) General workflow for preparing and analyzing unstained cell images using deep learning, including image generation, labeling, dataset preparation, model training, and performance evaluation. (B) Strategy for model selection and validation, starting with training and optimization on baseline datasets, followed by evaluation on test and unseen datasets to ensure generalizability

## Results

To investigate the use of DL to detect morphological differences in unstained cells, we focused on suspension cell lines cultured with the semi-automated AICE3 device with integrated image generation. Overviews of all created baseline and application data sets are shown in Tables 1 & 2.

### Single-class cell line detection

To investigate the ability of AI to detect unstained cells, we first investigated the human chronic myeloid leukemia cell line K562. In order to build a robust dataset, images with cells off focus, debris, bubbles and background only were included as negative controls (22 images). Ultimately, the cell detection dataset consisted of 436 images with 15.5 annotations (range 0-54) per image on average and 6,764 annotations in total (Table 1).

**Table 1.**
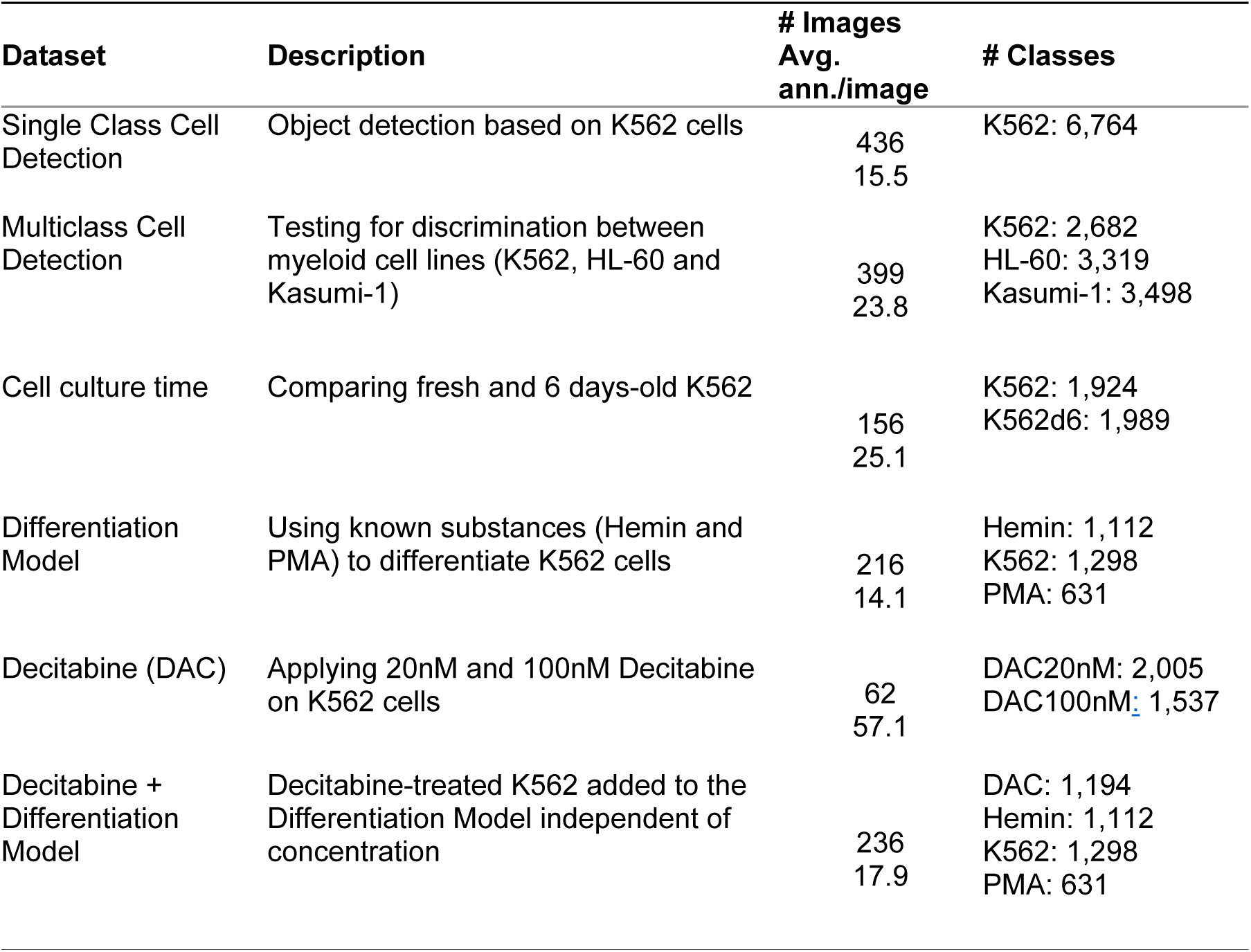
Overview of baseline datasets including name, description, abbreviation, and information per dataset including the number of images (# Images), average of annotations per image (Avg. ann./image), and the number of instances of corresponding classes (# classes)

| **Dataset** | **Description** | **# Images**<br>**Avg.**<br>**ann./image** | **# Classes** |
| --- | --- | --- | --- |
| Single Class Cell Detection | Object detection based on K562 cells | 436<br>15.5 | K562: 6,764 |
| Multiclass Cell Detection | Testing for discrimination between myeloid cell lines (K562, HL-60 and Kasumi-1) | 399<br>23.8 | K562: 2,682<br>HL-60: 3,319<br>Kasumi-1: 3,498 |
| Cell culture time | Comparing fresh and 6 days-old K562 | 156<br>25.1 | K562: 1,924<br>K562d6: 1,989 |
| Differentiation Model | Using known substances (Hemin and PMA) to differentiate K562 cells | 216<br>14.1 | Hemin: 1,112<br>K562: 1,298<br>PMA: 631 |
| Decitabine (DAC) | Applying 20nM and 100nM Decitabine on K562 cells | 62<br>57.1 | DAC20nM: 2,005<br>DAC100nM: 1,537 |
| Decitabine + Differentiation Model | Decitabine-treated K562 added to the Differentiation Model independent of concentration | 236<br>17.9 | DAC: 1,194<br>Hemin: 1,112<br>K562: 1,298<br>PMA: 631 |

**Table 2.** Overview of the validation datasets including name, description, abbreviation, and information about number of images for three datasets used for further validation.

| Abbr. | Dataset | Description | Number of images |
| --- | --- | --- | --- |
| LM | Labmaite K562 dataset | Other passage of K562 cells on a similar AICE system in Freiburg | 75 images |
| Mixture | Mixture dataset | Mixing cells with different substance treatment in a 50:50 ratio | 128 images |
| GC | Growth curve dataset | Two runs of K562 over nearly eight days in AICE system. Images were taken every two hours | 716 images |

All YOLOv8 detection models (s, n, m, l) achieved similar metrics. However, model size correlated with longer training times. Thus, in balance of speed and accuracy, the YOLOv8-s was chosen for further analysis. Object detection in this test set correctly identified 570 true positive K562 cells (95,6%) and only 26 were mislabeled - 15 as false positive/ type I error (mostly misidentified bubbles or debris) and 11 as false negative/ type II error (cells out of focus) (Figure 2B). Precision and recall/ sensitivity were over 97% (Figure 2C). Detailed metrics are shown in the supplements (Supplemental Table 2). Next, we transferred the robustness and reproducibility of the models to another laboratory, but with the same cultivation system. Therefore, K562 cells were cultured and imaged on a separate AICE3 device in Freiburg. This application dataset contained 75 images with variating cell densities. Although the precision was slightly lower due to misinterpreted background, no cells were missed (no false negatives) (Supplemental Figure 3). These results prove the reliability and accuracy of the model to detect unstained cells.

**Figure 2:**
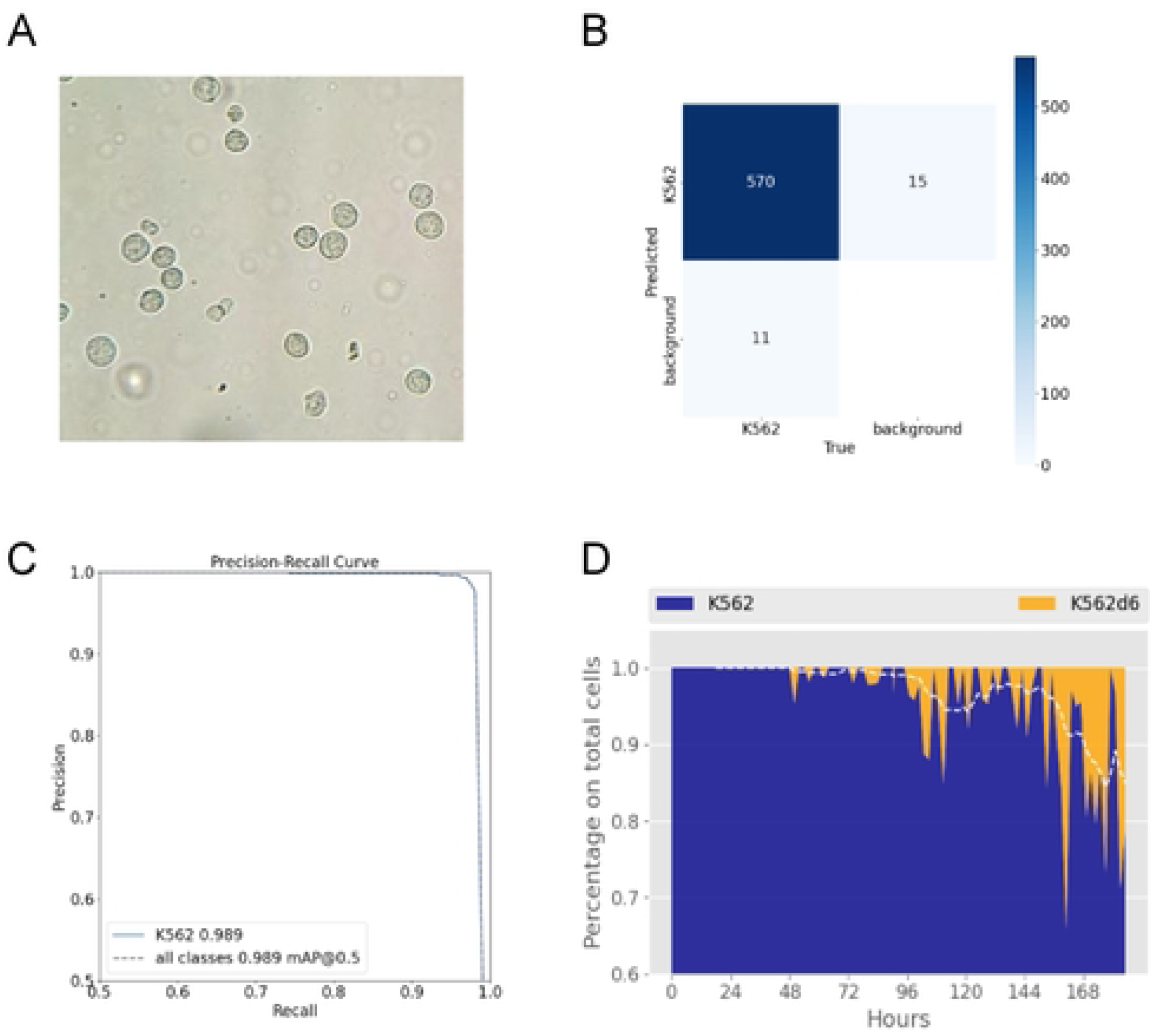
Performance evaluation and application of a deep learning model for cell classification. (A) Exemplary microscopic picture of K562 cells generated in the AICE3 device. (B) Confusion matrix showing the classification performance of the K562 model on the test set distinguishing K562 and background. Higher numbers are represented by darker shades of blue. (C) Precision-recall curve with the average precision (AP) for the K562 class and the overall mean average precision (mAP) at an IoU threshold of 0.5. (D) Stacked area plot showing the temporal distribution of K562 and K562d6 cells over a 8-day period (x axis), represented as a percentage of the total cell population (y axis). The moving average is shown as white dashed line.

Based on this dataset, we further tested the usability of the trained model for a real-time approach. Therefore, performances on a GPU and on two types of CPU environments (a Kubernetes/Kubeflow deployment and a local machine) were compared. On the GPU, prediction took 5.4 ms and 0.9 ms for pre- and post-processing steps respectively and 33.3 ms on average for 75 images (over 370 cells). Kubeflow CPU required 14.1 ms, 2468.9 ms, and 38.4 ms for preprocessing, average prediction and postprocessing, respectively, and the local CPU 16.3 ms, 1301.9 ms, and 1.5 ms. Therefore, GPU-, rather than CPU-, based analysis might be better suitable for real-time detection.

### Detection of longitudinal morphological changes

Variation in culture conditions over time associated with subtle morphological changes are expected to occur in prolonged culture. In order to discriminate these effects from substance-specific effects in a longitudinal observation of a drug testing setting, we investigated the impact of culturing time by reassessing and comparing K562 cells over a time span of six days. Firstly, K562 cells were kept in classic cell culture for six days (without splitting or media change) before image generation and were classified as K562d6. In total, the dataset comprises 156 images including 3,913 annotations with a mean of 25.1 cells per image (range 0-79). Five images show background only. A subset of the previous dataset with normal K562 was used as baseline reference. Here, the YOLOv8-n model achieved high precision and sensitivity (>95%), as well as ∼99% specificity, in distinguishing fresh K562 versus K562d6. The observed high mean average precision (mAP) scores indicate accurate cell framing (Supplemental Table 2).

Subsequently, to test the model in a dynamic longitudinal setup, K562 cells were kept for over seven days (190 hours) in the AICE3 device to continuously monitor cell counts and changes over time. Images were generated in a two-hour interval, resulting in 380 images. Over time, a gradual increase in the proportion of detected K562d6 cells can be observed reflecting the natural cell culture time associated changes of the culture (Figure 2D). These findings highlight the capability of an AI-based model to detect very subtle morphological changes in unstained cells not self-evident to the human eye and enables correction of culture time-associated effects.

### Multiclass leukemia cell line discrimination

Next, we added other morphologically highly similar myeloid cell lines to the model in a multi-class setup by including human HL-60 and Kasumi-1 cells (both acute myeloid leukemia cell lines). Aiming for a balanced dataset, only a subset of the K562 data was used considering both, number of images and cell density per image. In total the data set consisted of 399 images with a mean average of 23.8 annotations (range 0-139) per image resulting in 2,682, 3,319 and 3,498 annotations for K562, HL-60 and Kasumi-1, respectively.

Here, the model YOLOv8-m performed best. High mAP values (mAP@0.5 98.3%; mAP@0.5:0.95 74.3% (mAP across IoU thresholds 0.5 to 0.95)) indicate high overlap between annotated ground truth and the models bounding box output and classification prediction. A very high rate of true positive objects for all classes was found. Overall, misinterpretations were rare and mostly with background (cells or bubbles off focus). Solely in four instances K562 cells were confused for HL60 cells. Average sensitivity and specificity were above 97%, and precision was 94.6% (Figure 3 A,B). Detailed metrics are given in the Supplemental Table 1.

**Figure 3:**
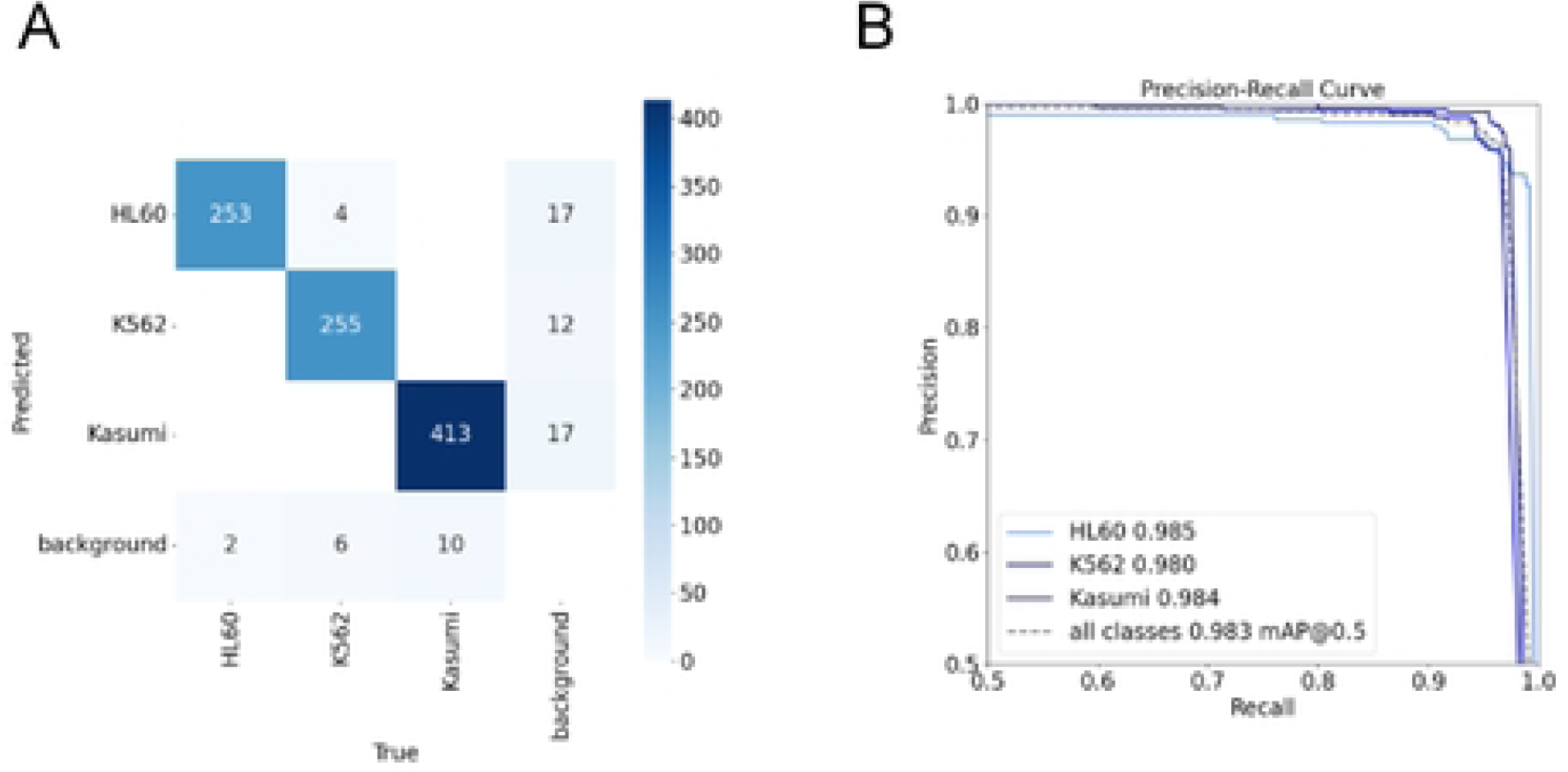
Performance evaluation of the deep learning model for myeloid cells. (A) Confusion matrix of predicted (y axis) versus true classes (x axis) for the myeloid cell lines HL-60, K562, and Kasumi-1. Higher numbers are represented by darker shades of blue. Correct classification is shown along the diagonal. Background classes holds cells that were not detected at all. (B) Precision-recall curves with the average precisions (AP) for HL-60 (light blue), K562 (mid blue), and Kasumi-1 (dark blue). The overall mean average precision (mAP) at an IoU threshold of 0.5 is shown as grey dashed line.

These results demonstrate that the trained model can precisely distinguish between three morphologically highly similar leukemia cell lines based on unstained microscopic images.

### Applying the cell line recognition model to a cellular differentiation model

We first focused on a previously described K562-based differentiation model (Figure 4 A) (13). K562 cells were treated with hemin and PMA for erythroid and megakaryocytic differentiation, respectively. In this model, not all cells fully transition into differentiation, but rather differentiate more or less. Cell morphologies suggest differences in cell size, opacity, and granularity between these classes (Figure 4B). This Differentiation Model dataset comprised 216 images with a total of 3,041 annotations (14.1 annotations per image on average) across the classes of hemin-treated, PMA-treated, and untreated K562 cells. In three instances, K562 cells were identified as PMA-treated cells and 15 annotations were originally background (debris or bubbles) but counted as cells. However, YOLOv8-s discriminated between the different classes with a very high precision, sensitivity, and specificity of approximately 95% (Supplemental Table 1). Therefore, the trained model was able to capture treatment specific morphological differences and reliably distinguish untreated K562 from Hemin and PMA-treated cells with great confidence according to the ground truth annotations.

**Figure 4:**
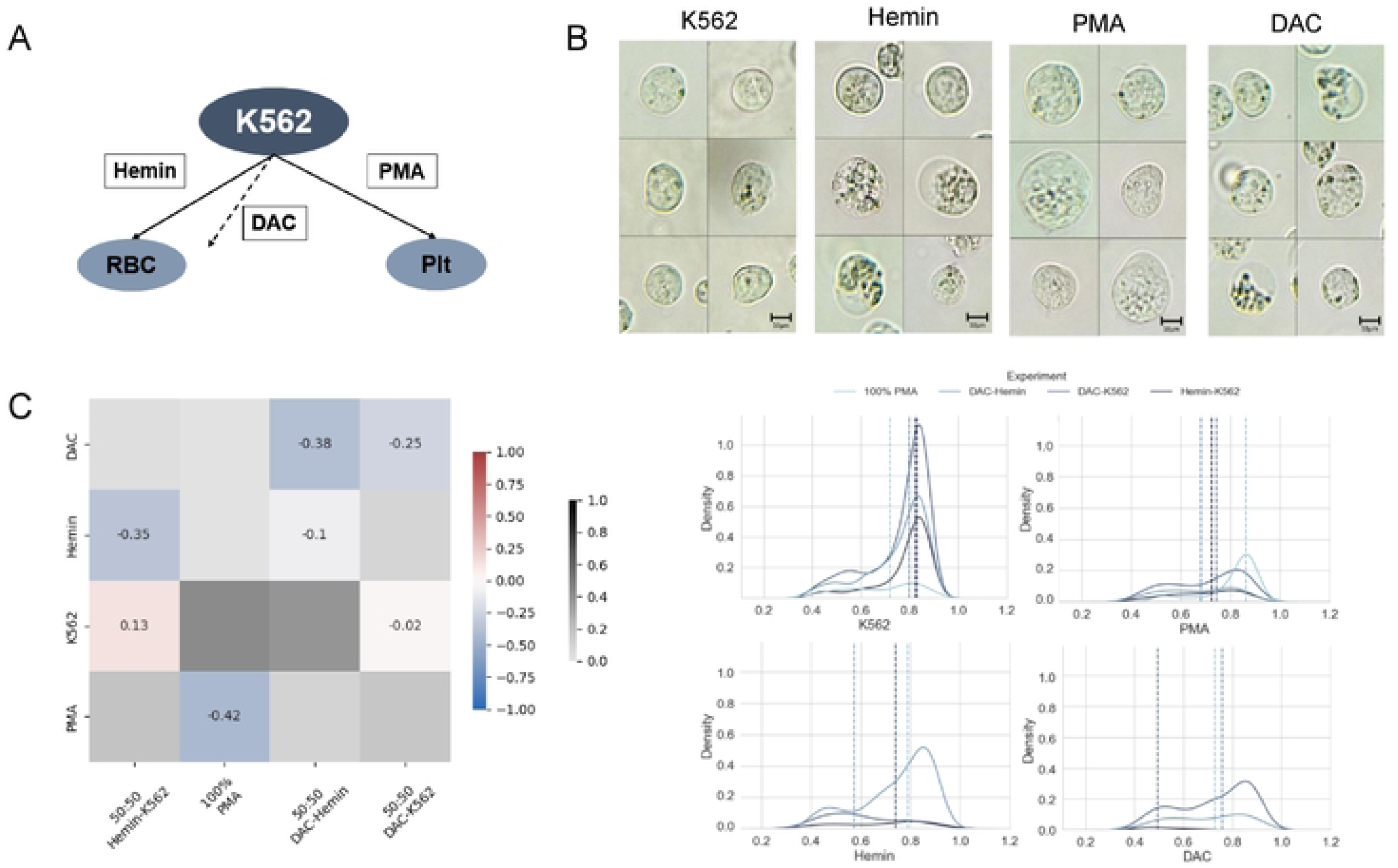
Schematic overview and performance evaluation of K562 cell differentiation upon treatment. (A) Model of differentiation from K562 cells towards red blood cells (RBC) under Hemin addition, towards platelets (Plt) under PMA, and under Decitabine (DAC) treatment towards RBC alike cells. (B) Representative brightfield images of untreated K562 cells and cells treated with Hemin, PMA, or DAC. Scale bars represent 10 μm. (C) Confusion matrix for validating mixures (x axis) on the drug testing model (y axis: K562 + PMA (PMA), (untreated) K562, K562 + Hemin (Hemin), and K562 + Decitabine (DAC)). Overestimation of a class by the model is shown in red color shades, underestimation corresponds to blue shades. Shades of grey represent a detection by the model although the class was not included in the experiment mixture. (D) Density plots (y axis) of the probability (x axis) for each class (K562, PMA, Hemin, DAC) showing the detection of the corresponding class in the experiments (lines: 100% PMA (light blue), DAC-Hemin (mid blue), DAC-K562 (blue), Hemin-K562 (dark blue). For each experiment, the mean is provided as vertical dashed line.

### Applicability of the cell differentiation model to a drug testing setup

Next, we used the hypomethylating substance Decitabine (Figure 4A). In patients with acute myeloid leukemia (AML), this agent induces fetal hemoglobin expression providing a useful dynamic biomarker of outcome. Additionally, Deciatbine has been shown to induce erythroid rather than megakaryocytic differentiation *in vitro* (*13*). Here, we introduced Decitabine-treated K562 cells (DAC) as a novel class to the training model.

First, we tested if object detection algorithms can recognize different substance concentrations. This dataset consists of 2,005 and 1,537 annotations of cells treated with 20 nM and 100 nM Decitabine, respectively. No tested model was able to accurately distinguish between the two concentrations applied (Supplemental Table 2). Based on these findings, both classes (20 and 100 nM) were merged into a single class “DAC” and 20 additional images with 1,194 annotations were included to the data set.

This new formed class DAC was included to the model of differentiated cells resulting in four distinct classes: untreated K562, Hemin, PMA, and DAC. Here, YOLOv8-m showed high precision, sensitivity and specificity, as well as high mAP scores (Supplemental Table 2). For validation, we included predefined mixing ratios of the four classes (Figure 4C, Figure 4D). For instance, DAC cells were exclusively detected in experiments in which they were included (DAC-Hemin / DAC-K562). When DAC and hemin cells were mixed (50:50), the hemin cells were identified with high precision and only slightly underestimated (-10%), while the DAC cells (-38%) were mostly misinterpreted as untreated K562 cells. This observation is similar in the PMA experiment, in which a high proportion was also incorrectly classified as untreated K562. In the 50:50 mixture of DAC and untreated cells, treated cells were underestimated by a quarter. However, untreated K562 cells were almost perfectly classified (98%).

### Morphological insights

Finally, we sought to add interpretability to the AI model’s strong performance. Therefore, we used the visualization technique Eigen-CAM to generate heatmaps highlighting regions that contribute to the decision making on the three leukemia cell lines (Figure 5). We used a public Eigen-CAM library for YOLOv8 (REF Rigved) on randomly picked cell crops.

**Figure 5:**
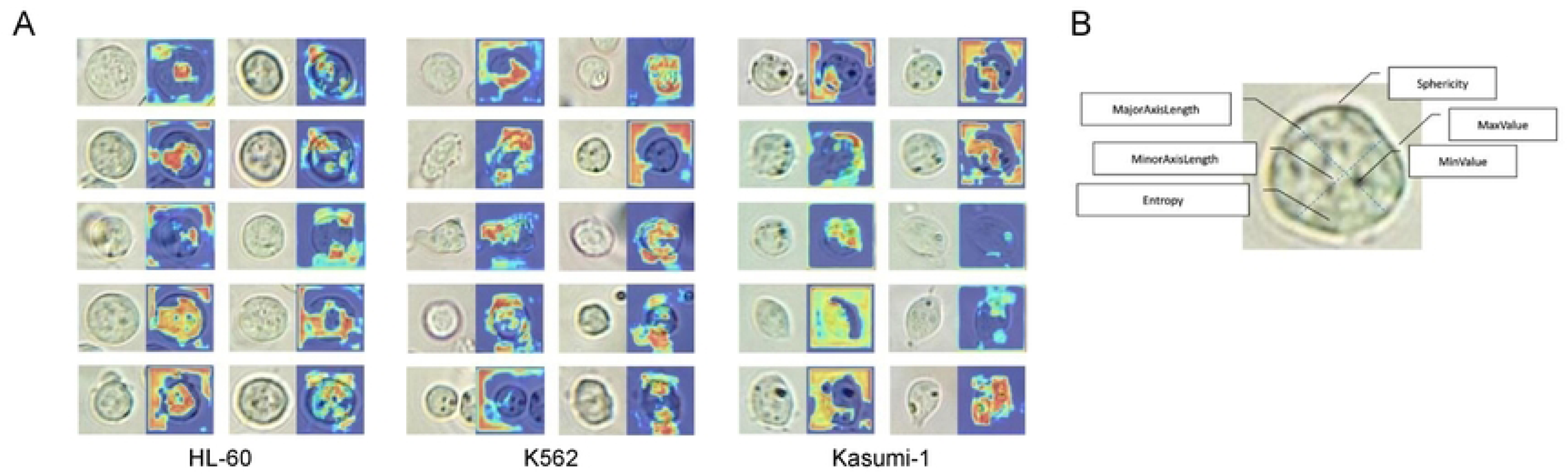
Visualization of cell-specific features contributing to deep learning classification. (A) Class activation maps (CAMs) overlaid on cell images for three different cell lines (HL-60, K562, Kasumi-1). Each pair shows the original cell image and the corresponding heatmap highlighting regions most influential for the model’s decision-making. (B) Schematic representation of selected morphological and texture-based features extracted from individual cells, including axis lengths, entropy, sphericity, and intensity metrics.

As exemplified in Figure 5A, Eigen-CAM was not always fully conclusive as also background was used for classification.

Beyond the DL–based approach, we also explored classical radiomic feature extraction (via RedTell) to compare conventional shape/texture descriptors with the model’s performance. This analysis addresses whether classification could be achieved using traditional features solely. In total, 74 features were extracted for each cell (exemplarily shown in Figure 5B). The Orthogonal Matching Pursuit (OMP) algorithm was used in order to select the best matching basis vectors from combinations of characteristics in an iterative manner. This approach was tested on the same datasets with the three cell lines, resulting in a binary classification for each cell line. An overview of the selected variables in the identified order is provided for each cell line in the Supplemental Table 3. For HL-60 cells the model focused primarly on the intensity of the energy, the RMS of the intensity, solidity and elongation, as well as gray level run length matrix (glrlm) variance and the perimeter surface ratio. Based on these features, 71% of HL-60 cells were classified successfully. For K562, the perimeter and three gray level emphasis features were used for the binary model. On the test data, the model achieved 82% accuracy. Eight variables (LRLGLE, minimum intensity, gray level variance, diameter, SRLGLE, maximum intensity, LGLRE, lmc12) were highlighted in order to classify Kasumi-1 cells. As a result, 78% of these cells were distinguished correctly. The corresponding Precision-Recall-curves for all classes are shown in Figure 6. Overall, the extracted features provide an explanation of the leukemia cell lines to some degree by using OMP. However, none of the explainable models depicted comparable high results as the AI-based approach.

**Figure 6:**
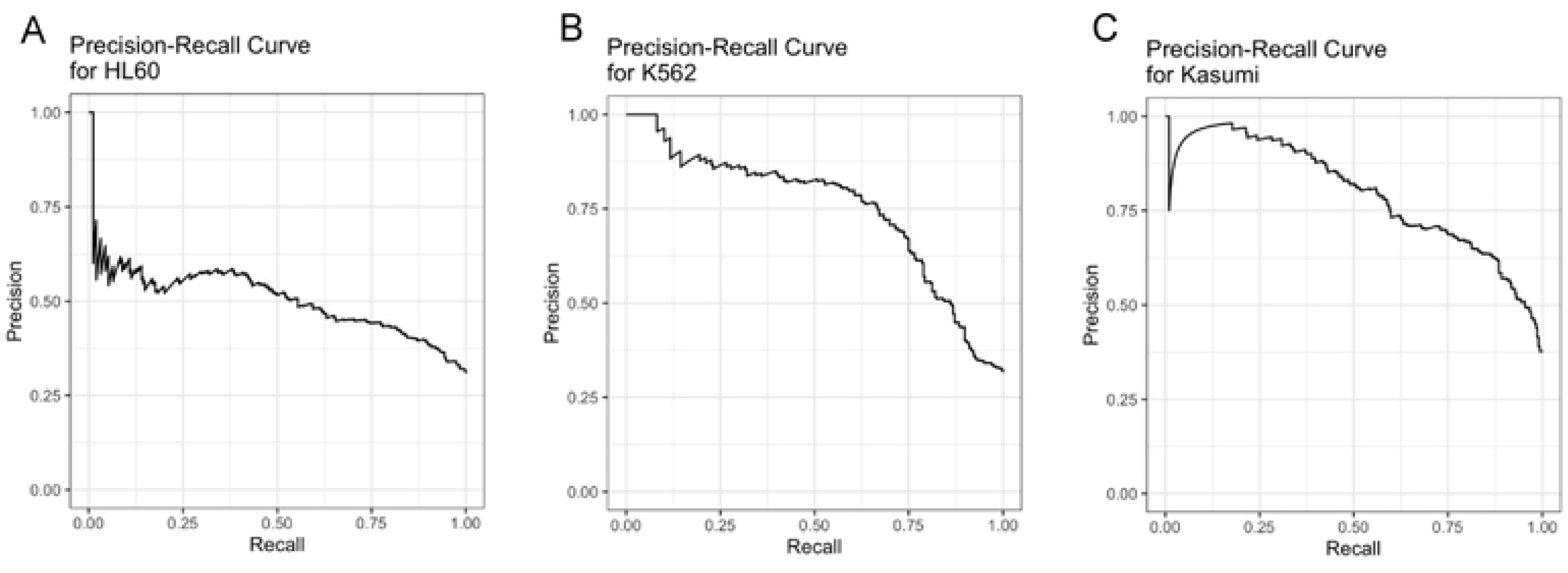
Precision-Recall curves for individual cell lines. (A–C) Precision-recall performance for the classification of HL-60 (A), K562 (B), and Kasumi-1 (C) cells using Orthogonal Matching Pursuit (OMP).

## Discussion

*In vitro* drug testing is an emerging tool in cancer research. It enables high-throughput screening in a controlled environment and supports personalized medicine by linking therapeutic responses to individual molecular/ genetic profiles. However, conventional staining-based measurement methods for assessing cellular responses have limitations, such as sample heterogeneity, potential effects on cell viability and often a lack of sequential measurement capabilities to assess dynamics. Image-based analysis of unstained cells offers a promising alternative, enabling real-time, non-invasive monitoring. Advances in DL and automated microscopy have facilitated high-throughput and unbiased cell classification. However, automated phenotyping of subtle morphological changes associated with drug response remains challenging. This study explores the feasibility and efficacy of image-based analysis on live, unstained and unfixed leukemia cell lines to enable longitudinal, high-throughput real-time screening.

Detection of unstained cells has been successfully demonstrated in different setups (17–19). However, these studies focused on cell groups with distinct morphological characteristics and phenotypic features. Therefore, we focused on the potential of deep-learning models in detecting and classifying unstained cells with highly homogeneous cell morphologies. AML is a case in which individualized treatment options and drug testing are starting to pave their way into modern treatment procedures (20). In addition, AML cell lines, which generally do not grow adhesively, enable largely automated cultivation and testing of drugs in a closed in vitro system.

Our trained AI models successfully identified unstained leukemia cells and, although background was challenging occasionally, it could discriminate against non-cellular components. Additionally, it recognized and classified similar cell lines with only subtle morphological variations with high accuracy. Also, detection speed is crucial for real-time assessment. In general, DL-based models need large GPU-based systems for fast execution and currently available GPU power allows for realtime assessment. However, we showed that a local CPU device can also run our trained model rapidly. This enables analysis on basic hardware without specialized infrastructure.

In summary, these results lay the foundation for complex experiments with a straightforward, accessible, and fast readout, enabling measurement of subtle changes (e.g. specific responses to certain substances) in a drug screening setup in near real-time.

By eliminating pre-processing steps like staining and fixation, cytotoxic effects are avoided. This permits to keep the cells in a closed system, fostering a feedback loop that enhances experimental efficiency and allowing longitudinal experiments (21). However, cells in cultivation carry out physiological processes naturally, such as metabolizing and consuming media over time. Additionally, cells undergo nutrient depletion and waste accumulation (22).

These effects can lead to subtle but significant morphological changes which may play a critical role in longitudinal drug testing experiments (23). Hence, we tested if an AI model is capable to perceive any changes over time solely based on unstained cell morphology. In longitudinal experiments, our AI model tracked a morphological drift with increasing culture time. Thus, the model successfully detected time-dependent variations and recognized subtle cellular changes not obvious to the human-eye. Particularly in the context of longitudinal drug testing, our results show the potential to discriminate between substance-specific and culture-time associated effects – a key element for assessing drug efficacy and cellular adaptation in a controlled environment.

We further explored the ability of our models to detect subtle morphological changes induced either by time in culture or by specific treatments. Cell culture time-related drift in morphology was detected over the course of continuous culture, emphasizing the need to account for such effects when interpreting drug responses. When applied to a drug testing context, the model accurately differentiated between K562 cells undergoing erythroid or megakaryocytic differentiation, as well as those treated with Decitabine. Interestingly, the classifier appeared to reflect a gradient of differentiation states rather than binary transitions, suggesting the model is sensitive to intermediate phenotypes—an important consideration for tracking dynamic cellular responses.

However, one of the key challenges in AI applications for biomedical imaging is the understanding how models make their decisions. While visualization techniques such as Eigen-CAM offered some insights into feature prioritization, they were not wholly informative. Although focus on submembranous regions and cytoplasmatic areas was recognizable in some instances, background was also highlighted to a great extent and cell membranes were barely considered. In contrast, classical radiomics-based feature extraction offered more transparent, although less accurate, models. This contrast underscores a central trade-off in AI model development: the balance between performance and explainability.

In summary, the findings of this study represent a relevant step towards fully automated drug testing platforms based on a simple, reliable and fast readout. Compared to human-based evaluation, automation enhances stability and reduces variability, making the analysis more reliable and scalable. By ommitting staining and fixation, the time required for sample preparation is minimized and allows real-time assessments and continuous monitoring of living cells. The adaptability of AI to various time points in a feedback-loop opens new possibilities for studying cellular dynamics and therapeutic responses. This approach could enable the fast selection of effective treatments, potentially surpassing traditional transcriptomic or proteomic analyses in speed and efficiency. Our study highlights the transformative potential of computer vision models in drug screening, offering a novel perspective where AI-driven morphology-based assessments complement and enhance conventional methodologies. Furthermore, integrating AI-driven morphological analysis with algorithms designed to intelligently select and combine drugs could revolutionize the way to faster, more effective, and highly individualized treatment adaptation and selection.

However, several limitations must be acknowledged. Our system currently relies pimarily on bounding box annotations. While segmentation can be introduced in future iterations via smart polygon tools, this will require additional computational resources and validation.

With the automated conversion of bounding boxes into smart polygons via Roboflow, segmentation algorithms have now become accessible for future analyses. Next, our study was limited to two object detection algorithms, YOLOv8 and RT-DETR, which were chosen based on their strong performance in real-time detection tasks (16). While these architectures provide state-of-the-art results, further comparisons with alternative models could refine and improve our approach. A notable limitation of our study is the absence of a definitive ground truth for our experiments. Although image recognition allows for relative quantifications and pattern identification, we cannot fully validate the detection of e.g. erythroid and megakaryocytic differentiation at the single-cell level, as we havenot incorporated confirmatory techniques such as hemoglobin staining or immunophenotyping of differentiation markers of the identical cells. Furthermore, our experiments were restricted to a limited number of suspension cell lines and tested substances. Adding parallelization to our setup could significantly increase data generation. Another practical limitation is that modifications to the imaging system, such as changes in resolution, lighting or hardware, would require re-training the models to ensure accuracy and performance. This challenges the generalizability of the model across differend imaging platforms. The variations in image acquisition can limit the transferability of our model to other datasets so that its applicability outside of the current setup requires further validation and adaptation. Finally, the extent to which our findings apply to other biological settings, such as adherent cells, primary patient cells or co-culture scenarios, remains uncertain and requires further validation. Addressing this in future studies would be essential for a broader applicability of our approach, particularly for drug testing across diverse cellular environments.

Future research will focus on evaluating the applicability of our system across a broader range of technical (e.g. integrating methods with other assays) and biological (e.g. additional cell types) contexts. Additionally, we aim to shift from primarily detection-based analyses to more advanced segmentation-based methodologies; further refining the precision and depth of our assessments. A particularly promising application of this technology is its potential integration into feedback-loop systems for personalized medicine. AI-driven models could continuously monitor a patient’s response to treatment, adjusting medication selection and dosage in near-real-time based on image-based analyses and patient-specific data. This iterative process would enable highly individualized therapeutic strategies, enhancing treatment efficacy and improving patient outcomes. Ultimately, our approach could contribute to a more precise and adaptive framework for clinical decision-making, advancing the field of personalized medicine.

## List of abbreviations (in order of appearance)

DL: deep learning
CNN: convolutional neural network
DAC: decitabine
RT-DETR: real-time detection transformer
AML: acute myeloid leukemia
AI: artificial intelligence
mAP: mean average precision
OMP: orthogonal matching pursuit
PCA: principal component analysis

## Authorś Contributions

Conception and Design: RC, MT

Provision of study materials: JB

Collection and assembly of data: TM, MK, DR, EM, AS

Data analysis and interpretation: KH, TM, AR, SS

Manuscript writing: KH, TM, RC

Visualization: KH

Final approval of manuscript: all authors

## Acknowledgements

We would like to thank all staff of the BioThera-Roland-Mertelsmann Foundation and LabMAite.

